# MiRGraph: A hybrid deep learning approach to identify microRNA-target interactions by integrating heterogeneous regulatory network and genomic sequences

**DOI:** 10.1101/2023.11.04.565620

**Authors:** Pei Liu, Ying Liu, Jiawei Luo, Yue Li

**Affiliations:** College of Computer Science and Electronic Engineering, Hunan University, Changsha, Hunan, China; School of Computer Science, McGill University, Montreal,Quebec,Canada

**Keywords:** MiRNA-Target Interaction, Heterogeneous Graph Transformer, Deep learning, Transformer-based CNN, Gene regulatory networks, Genomic Sequences

## Abstract

MicroRNAs (miRNAs) mediates gene expression regulation by targeting specific messenger RNAs (mRNAs) in the cytoplasm. They can function as both tumor suppressors and oncogenes depending on the specific miRNA and its target genes. Detecting miRNA-target interactions (MTIs) is critical for unraveling the complex mechanisms of gene regulation and promising towards RNA therapy for cancer. There is currently a lack of MTIs prediction methods that simultaneously perform feature learning from heterogeneous gene regulatory network (GRN) and genomic sequences. To improve the prediction performance of MTIs, we present a novel transformer-based multiview feature learning method – MiRGraph, which consists of two main modules for learning the sequence-based and GRN-based feature embedding. For the former, we utilize the mature miRNA sequences and the complete 3’UTR sequence of the target mRNAs to encode sequence features using a hybrid transformer and convolutional neural network (CNN) (TransCNN) architecture. For the latter, we utilize a heterogeneous graph transformer (HGT) module to extract the relational and structural information from the GRN consisting of miRNA-miRNA, gene-gene and miRNA-target interactions. The TransCNN and HGT modules can be learned end-to-end to predict experimentally validated MTIs from MiRTarBase. MiRGraph outperforms existing methods in not only recapitulating the true MTIs but also in predicting strength of the MTIs based on the *in-vitro* measurements of miRNA transfections. In a case study on breast cancer, we identified plausible target genes of an oncomir.

## I. Introduction

MicroRNAs (miRNAs) are endogenous, non-coding, and single-stranded small RNA molecules composed of approximately 22 ribonucleotides. They serve as the post-transcriptional mediators by binding to the 3’ UTRs of the target mRNAs, thereby inducing mRNAs degradation or inhibiting mRNA translation [14]. MiRNA are implicated in almost all biological processes in human cells [2; 6]. Their dysregulation can affect gene regulatory programs ultimately leading to diseases including cancers [3]. Therefore, accurate prediction of miRNA-target interactions (MTIs) is of great significance for revealing the complex diseases regulatory mechanism and for designing miRNA-based therapy [9].

While traditional experimental methods were developed to detect MTIs [13], they are limited to a specific miRNA. Highthroughput sequencing of RNAs isolated by cross-linking immunoprecipitation (HITS-CLIP) or crosslinking, ligation, and sequencing of hybrids (CLASH) are costly and biased towards strongly bound MTIs [27]. Therefore, computational MTI prediction methods are cost-effective ways to prioritize candidate MTIs for experimental validation.

The computational algorithms of MTI predictions can be divided into two categories: rule-based methods and machine learning (ML)-based methods. The former utilize hand-crafted features. For instance, PITA [12] leveraged site accessibility to calculate thermodynamic stability-based scores. miRanda [5] utilized the sequence complementarity to assess the likelihood of MTI. TargetScan [1] prioritized the miRNA-candidate target site (CTS) using seed region matching rule. Although these rule-based methods provide putative MTIs, they depend on the design of hand-crafted feature engineering strategies, resulting in inconsistent predictions and high false positive rates.

ML-based methods train a supervised classification method to predict MTIs using a set of features. For instance, DeepTarget [15] adopted four types of sequence-based seed-matching features along with learned sequence-based features from LSTM-based autoencoder to predict CTSs. miRAW [22] applied multi-layer perceptron (MLP) to predict binding site accessibility energies. TargetNet [20] utilized a ResNet-based deep learning framework with a miRNA–CTS sequence encoding scheme to incorporate alignment of extended seed-region and mRNA sequences. Although these methods use CTS to narrow the search space, they rely solely on RNA sequence information and fail to incorporate the miRNA-mediated gene regulatory network (GRN) information.

Meanwhile, with the expansion of CTS databases like mirRTarBase [11] and TargetScan [1], it has become feasible to use graph neural networks to extract relational information between miRNA and mRNA species. MRMTI [18] utilized the multi-relational graph convolutional module and the Bi-LSTM module to incorporate both network topology and sequential information. SRG-Vote [28] leveraged Doc2Vec and Graph Convolutional Network (GCN) to separately generate sequential and geometrical embeddings from molecular sequences and relational network. Both methods operate on k-mers embeddings (i.e., k = 3 nucleotides). Specifically, they split a gene sequence into k-mer segments treated as “words” and then maps them to real-valued embeddings via a pre-trained word2vec-based model. In addition to the above shortcomings, there is currently a lack of MTI prediction methods for extracting node features from heterogeneous networks by modeling structural and relational data. Moreover, while considering edge and node information simultaneously in heterogeneous graph networks has proven to be beneficial [23; 19], their applications on MTI prediction remains unexplored.

In this study, we present a novel transformer-based deep learning framework named MiRGraph, which learn the MTI from a comprehensive miRNA-mRNA network and mRNA+miRNA sequence information (Fig. 1). Our contributions are two-fold. As the first contribution, our method utilizes the entire raw sequences of mature miRNA and the 3’UTRs of target mRNAs to learn the sequence features. This is in contrast to most existing methods that rely on short k-mer to represent sequences. To efficiently learn features from the entire 3’UTR sequences, we design a hybrid CNN and transformer architecture (TransCNN). As the second contribution, we construct a comprehensive heterogeneous graph network consisting of miRNA-miRNA from miRNA family information, gene-gene interaction from protein-protein interaction (PPI) network and MTI from TargetScan [1] and miRTarbase [11]. To efficiently learn the relational and topological information from the heterogeneous interactome network, we utilize a heterogeneous graph transformer (HGT) module. The concatenated sequence-based and graph-based feature embeddings are then fed to an MLP layer followed by a bilinear transformation function to compute the MTI prediction scores. As groundtruth, we use the experimentally validated MTIs from miRTarBase for classification task and *invitro* log-fold-change (LFC) of gene expression upon miRNA transfections for regression task. Comprehensive experiments show that MiRGraph outperforms the state-of-the-art methods in MTIs prediction tasks. In a case study, we use MiRGraph to prioritize *de novo* target genes of the oncomir *hsa-miR-122-5p* and *de novo* miRNAs associated with the oncogene *BRCA1* in breast cancer and find hits that are supported by existing literatures and databases.

**Fig. 1.**
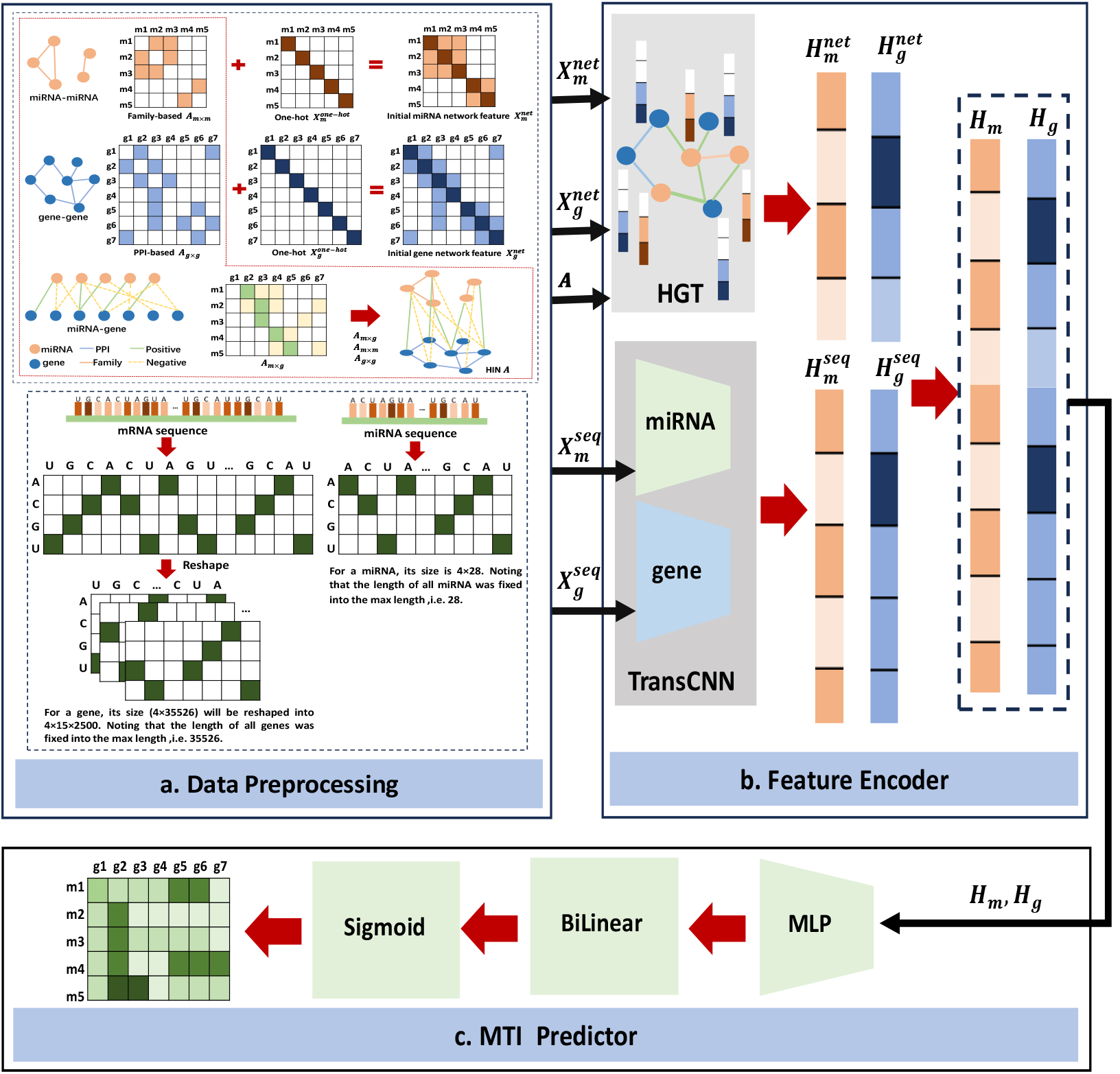
Overall framework of MiRGraph. **a**. HIN construction. We built a HIN *A* that comprises miRNA-miRNA, gene-gene and miRNA-gene interaction networks (*A*_*m×m*_, *A*_*g×g*_ and *A*_*m×g*_). And then we obtained the initial miRNA and gene network features 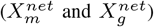by combining their one-hot features 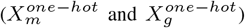and adjacency matrices of networks (*A*_*m×m*_ and *A*_*g×g*_). Also, we obtained one-hot encodings of the miRNA and gene sequences 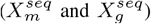. **b. Feature Encoder**. We first learn from the HIN and sequences feature embeddings of miRNAs 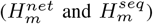 and genes 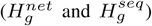, using ‘HGT’ and ‘TransCNN’ modules separately. We then concatenate them into the final feature embeddings of miRNAs and genes (*H*_*m*_ and *H*_*m*_). **c. MTI predictor**. We feed the learned features of miRNAs and genes (*H*_*m*_ and *H*_*m*_) into MLP. Finally, we use the ‘BiLinear’ and ‘Sigmoid’ functions to calculate their interaction scores.

## II. MiRGraph framework

The proposed framework MiRGraph mainly consists of three parts (Fig. 1): a. Heterogeneous information network (HIN) construction; b. Feature Encoder; c. MTI Predictor. Briefly, we first built a heterogeneous information network (HIN), and obtain the initial network and sequence one-hot encoding of miRNAs and genes (Fig. 1a). They are fed into feature encoder including two modules (i.e. HGT and TransCNN) to learn network and sequence representations separately (Fig. 1b). Finally, we feed the concatenated representation into the MTI predictor (Fig. 1c). Details of each step are described below.

### A. Data Preprocessing

#### Heterogeneous GRN Construction

The predicted miRNA-gene interactions network was obtained from TargetScan [1]. We used the experimentally validated MTI from miRTarbase [11] to label the TargetScan-predicted edges as positive edges and the rest as negative edges. The above procedure resulted in 323,370 positive edges and 666,932 negative edges between 2656 miRNAs and 18,454 genes. Considering that the negative edges of the network built on this basis are too large, we retained only the negative edges for each gene *g*, which involves the bottom 5 miRNAs with sequence most similar to the miRNA involved in the positive MTI. This reduces the negative samples and improves the specificity of our model to discriminate the ture MTIs. We obtained the gene-gene network from the PPI network with 13,098,934 edges between 18,454 genes from the STRING database [25]. The miRNA-miRNA interaction network with 16102 edges between 2656 miRNAs was built with miRNA family information [17]. The resulting HIN *A*_(*m*+*g*)*×*(*m*+*g*)_ ∈ [0, 1]_(2656+18454)*×*(2656+18454)_ consists of three networks, i.e., the miRNA-gene (*A*_*m×g*_ ∈ [0, 1]^2656*×*18454^), gene-gene (*A*_*g×g*_ ∈ [0, 1]^18454*×*18454^), and miRNA-miRNA (*A*_*m×m*_ ∈ [0, 1]^2656*×*2656^).

#### Feature Initialization

We converted the miRNA sequences into one-hot representations with the maximum length set to *k*_*m*_ = 28. We processed the target gene sequences in the same way with the maximum length set to *k*_*g*_ = 35, 526. Because the maximum length of the complete 3’UTR sequences of all genes is too long, each gene sequence was divided into 15 fixed-length shorter sequences. As a result, for all miRNAs and genes, we obtained their initial sequence features 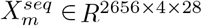and 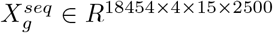. This is equivalent to dividing a long 2D image into 15 segments each having height of 4 and width of 2500.

To obtain the initial miRNA and gene network node features, we first constructed the diagonal matrix based on miRNAs and genes identities using one-hot encoding i.e. 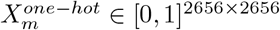and 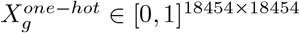. To enrich node features, we integrated the one-encoded node feature with the corresponding network matrices via elementwise addition: 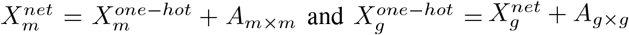(Fig. 1a).

### B. Feature Encoder

#### Sequence Feature Encoding Module

In order to fully capture the MTI-specific sequence elements in miRNA and gene sequences, we designed a novel sequence learning module TransCNN by combining the convolutional neural network (CNN) and the Transformer encoder (Fig. 2). Specifically, for gene 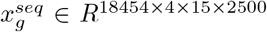, we devised a three-layer encoder namely *TransCNN gene* (Fig. 2a):

**Fig. 2.**
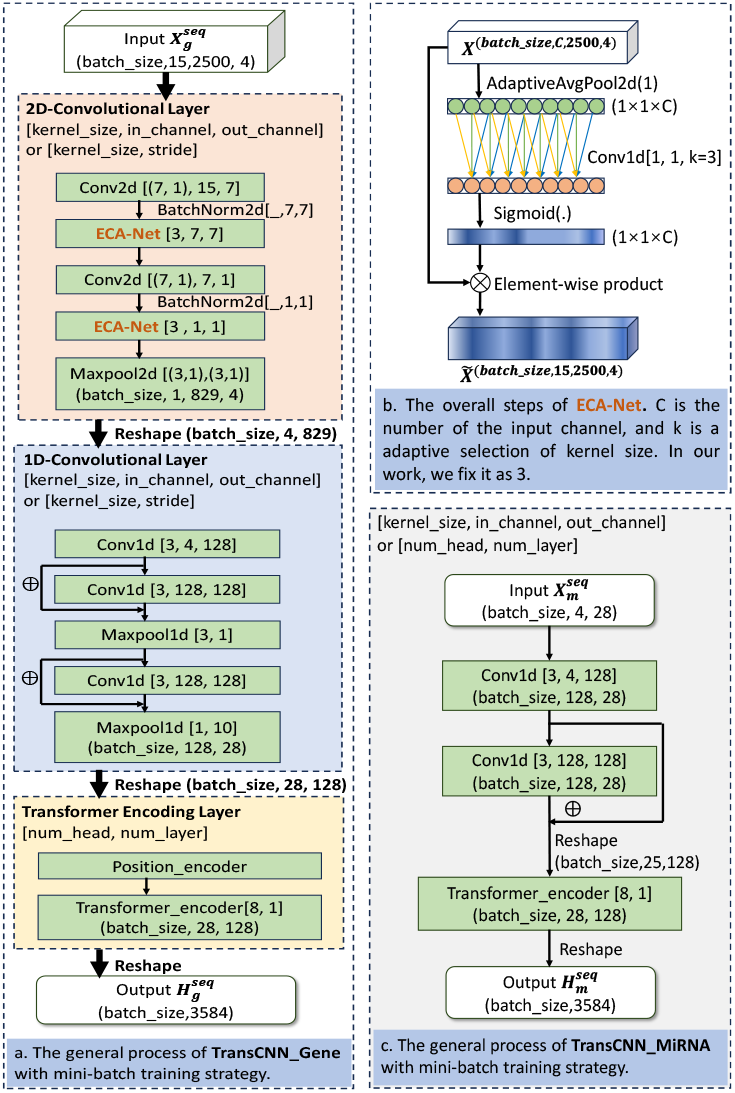
The TransCNN module. The diagram depicts the sequence encoder module as the part of the MiRGraph encoder (Fig. 1b). It is composed of two sub-modules namely TransCNN Gene and TransCNN MiRNA. **a**. TransCNN Gene. This sub-module consists of three layer types: ‘2D Convolutional layer’ used to learned cross-channel interaction information, ‘ResNet layer’ used to learned intra-channel specific information, and ‘Transformer encoding layer’ used to capture more fine-grained information. **b**. Detailed view of ECA-net. ECA-net is used to model local cross-channel attention values. **c. TransCNN MiRNA**. Due to short sequences of miRNA, this sub-module only uses residual connection after the 1D convolution layer *conv*1*d*(.), followed by transformer encoder.

- **2D convolutional layer**. Two layers of 2D convolutional layer *conv*2*d*(.) are used to integrate the 15 sequence segments of each gene. This is followed by a max-pooling layer *maxpool*2*d*(.) to reduce the sequence length. To effectively learn features, we treat the 15 sub-sequences as 15 channels and introduce a channel attention mechanism layer (ECA-Net) [26] between the two *conv*2*d*(.) layers. **ECA-Net** improves the expressiveness of the feature representation by a set of globally adaptive weights of the feature map of each channel. Specifically, it first dynamically determines the kernel size *k* after aggregating convolution features using global average pooling (GAP) without dimensionality reduction, and then performs 1D convolution followed by Sigmoid transformation to learn the channel attention. Finally, it calculates the elementwise product between channel attention and original feature to obtain the enhanced feature (Fig. 2b).
- **Residual connection layer**. Residual connection is adopted after the 1D convolution layer *conv*1*d*(.). A maxpooling layer *maxpool*1*d*(.) is applied after each residual connection to further reduce the sequence length.
- **Transformer layer**. One layer of transformer encoder *transformer encoder*(.) is adopted to learn global contextual information from the sequence feature map.

For the input of miRNAs 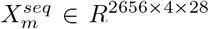, we utilize similar operations in steps 2 and 3 in *TransCNN gene* to learn sequence features of miRNAs (Fig. 2c). In order to integrate the two types of sequence information, we designed the convolution filters to match the dimensions between the final sequence feature embedding of miRNAs and genes, i.e., 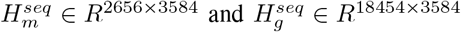.

#### Network Feature Encoding Module

Our HIN graph *A* = (*V, E, B, D*) comprises node *v ∈ V*, edge *e ∈ E*, and the corresponding node and edge types *B* and *D*, which can be mapped via functions *τ* (*v*) : *V* → *B* and *θ*(*e*) : *E* → *D*. For any edge *e* = (*s, t*) from source node *s* to target node *t*, its meta-relation is expressed as ⟨*τ* (*s*), *θ*(*e*), *τ* (*t*)⟩. We adapted the Heterogeneous Graph Transformer (HGT) model [10], which uses the meta-relations of the heterogeneous graphs to parameterize its weights for computing the heterogeneous mutual attention, messages, and propagation steps. HGT can automatically learn and extract the salient ‘meta-paths’ for different downstream tasks, even if the input to HGT is only one-hop edge without manually designed meta-paths. Specifically, it is composed of three key parts as follows.

#### Heterogeneous Mutual Attention

Given a target node *t* and a source node *s ∈ N* (*t*), where *N* (*t*) is the set of neighbor nodes of *t*, their mutual attention (*Attention*(*s, e, t*)) grounded by their meta relations ⟨*τ* (*s*), *θ*(*e*), *τ* (*t*)⟩ triplets is defined as the following equations:

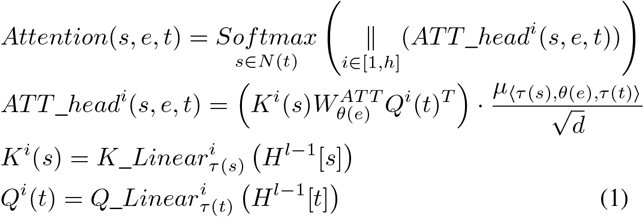

where ∥ (.) is the concatenation function. *ATT head*^*i*^(*s, e, t*) is the output of the *i*^*th*^ attention head between *s* and *t*. 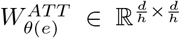is a edge-based matrix to learn different meta-relation features even between the same node type pairs, where *h* is the number of attention heads and ^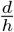^ is the vector dimension per head. (.)^*T*^ is the transposal function. *µ*_*<τ* (*s*),*θ*(*e*),*τ* (*t*)*>*_ is a prior tensor, serving as an adaptive scaling of attention to denote the significance for each meta-relation. *K*^*i*^(*s*) is the *i*^*th*^ Key vector of source node *s* and *Q*^*i*^(*t*) is the *i*^*th*^ Query vector of target node *t*, and they are separately mapped by the linear projection functions 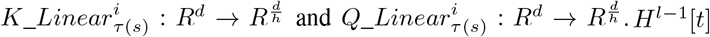are node embedding of nodes *t* and *s* at the *l* − 1^*th*^ layer, respectively.

#### Heterogeneous Message Passing

To pass information from source nodes to target nodes, the multi-head message for a pair of nodes *e* = (*s, t*) is calculated by:

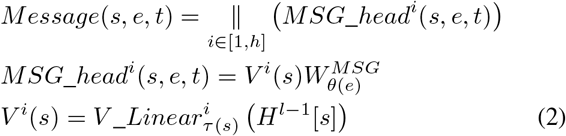

where *MSG head*^*i*^(*s, e, t*) represents the *i*^*th*^ head message between *s* and *t*. 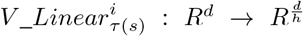is a linear projection function to map each source node *s* in the *i*^*th*^ head into a message vector *V* ^*i*^(*s*). 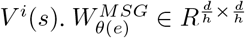is a edge-based matrix similar to 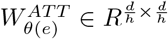.

#### Target-Specific Aggregation

To get the output *H*^*l*^[*t*] of the target node *t* after *l* layers, we first aggregate the heterogeneous multi-head attentions and messages from all source nodes *s* ∈ *N* (*t*) to target node *t*, and then use a linear projection function *A Linear*_*τ*(*t*)_ to update the embedding 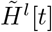followed by a residual connection:

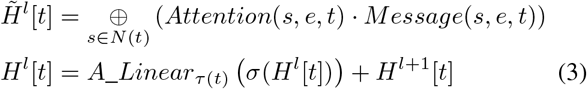

where *σ*(.) is an activation function. Through the above three steps, HGT can generate a highly contextualized representation *H* for each node in the heterogeneous graph. In summary, Fig. 3 depicts our HGT module, which outputs the network features of miRNAs and genes i.e., 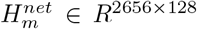and 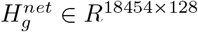.

**Fig. 3.**
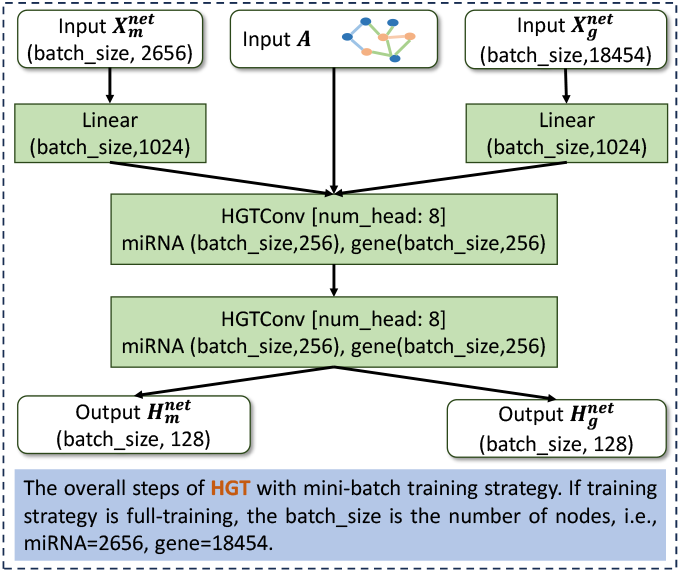
The HGT module. The diagram depicts the network feature encoder module as part of the MiRGraph encoder (Fig. 1b). Given a heterogeneous graph *A* with two node types *B* = *{miRNA, gene}* and three edge types *D* = *{′miRNA − miRNA*^*′*^,^*′*^ *miRNA − gene*^*′*^,^*′*^ *gene − gene*^*′*^*}*, we first feed the initial network features of miRNAs and genes 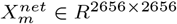and 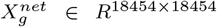to two separate linear layers *Linear*(.). The resulting node features along with the three adjacency matrices (i.e., *A*_*m× g*_ *∈* [0, 1]_2656*×*18454_, *A*_*g× g*_*∈* [0, 1]_18454*×*18454_ and *A*_*m× m*_*∈* [0, 1]^2656*×*2656^) are fed into the *HGTConv*(.) layer to learn the nuanced features of different types of nodes. After two layers of *HGTConv*(.), we obtain the final network feature representations of miRNAs and genes.

#### Feature Integration Module

We first concatenate the two types of node embeddings, and then apply the nonlinear activation function *GELU* (.) to obtain the final representation of miRNAs and genes, i.e. *H*_*m*_ ∈ *R*^2656*×*3712^ and *H*_*g*_ ∈ *R*^18454*×*3712^ (i.e., 3584 + 128 = 3712 (Fig. 1b):

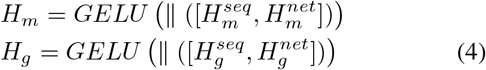

#### Predicting Score

Given the input of *H*_*m*_ and *H*_*g*_, we first use *MLP* (.) to separately map them onto the same low-dimensional space, and then use a bilinear layer followed by sigmoid function *Sigmoid*(.) to compute the MTI probability:

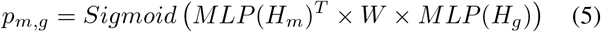

#### Loss Function

To account for the imbalance of small number of positive and large number of negative examples (i.e., true and false MTI edges), we use the focal loss function [16] to train the model:

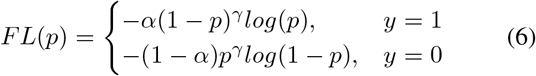

where *y* is the true label, *p* is the predicting score, *α >* 0 is a hyperparameter to balance the importance of positive/negative examples, and *γ >* 0 is a hyperparameter. By default, we set *α* = 0.65 and *γ* = 2.

## III. Experiments

### A. MTI prediction tasks

We train MiRGraph to predict candidate MTIs. We designed two scenarios: in scenario 1, we divided the dataset based on edges (i.e., MTIs). In details, we split the positive (miRTar-base+TargetScan) and negative edges (TargetScan only) into 90% and 10% for training and testing, then used 10% from the training set for validation; in scenario 2, we divided the dataset based on nodes (i.e., miRNAs) using the same splitting rule. In both scenarios, we used the complete HIN, which encompasses all 2,625 miRNAs and all 18,454 genes.

To further demonstrate the utility of MiRGraph, we used the log-fold-change (LFC) of miRNA interference (miRNAi) experiments followed by gene expression profiling by 9 microarrays for 9 miRNAs tested in HeLa cell line [7] and 2 RNA-seq experiments for 2 miRNAs (hsa-miR-1, hsa-miR-124-3p) tested in HEK293 cell line [21]. Following scenario 1, we randomly divided the LFC data into 90% and 10% for training and testing sets. We fine-tuned the miRGraph trained on miRTarBase by minimizing the regression smooth L1 loss, which is less sensitive to outliers than mean squared error loss.

### B. Baseline methods and Implementation details

We compared MiRGraph with several state-of-the-art methods, namely Relational Graph Convolution Network (RCGN) [23], MRMTI [18], HGT [10], HGT BiLSTM and TargetNet [22]. We used the original implementations from the above studies with default settings whenever available. When the code were not available, we implemented the method based on the description in the original paper using Pytorch (v 2.2.1) and PyTorch Geometric (v 2.5.0) as backend in Python (v 3.11.8). To apply them to MTI prediction task, we adopted the operator *RGCNConv* and *HGTConv*, implemented in *torch geometric*.*nn* package. MRMTI is an MTI prediction model using RCGN and Bi-LSTM to learn network and sequence information separately. HGT BiLSTM can be seen as a variant of MRMTI, which combines HGT and Bi-LSTM. Integrating RGCN and HGT with Bi-LSTM, we implemented MRMTI and HGT BiLSTM. Bi-LSTM was implemented by the operator *LSTM* in *torch*.*nn*. We leveraged Kaiming [8] to initialize the model parameters and AdamW as the optimizer for model training.

To efficiently learn from the large HIN graph (i.e., 18,454+2656 by 18,454+2656), we designed two strategies:

- **End-to-end transfer learning**. We first pre-trained TransCNN and HGT (i.e., separately training TransCNN+MLP and HGT+MLP) (Figure 1b). We then transferred the parameters of the trained HGT and TransCNN to MiRGraph (i.e., [TransCNN, HGT] + MLP) for an end-to-end fine-tuning (Fig. 1c).
- **Step-by-step training**. We first trained TransCNN and HGT separately. We fixed the HGT and TransCNN and only trained the prediction layer using the combined feature representations of miRNAs and genes from the two modules for final MTI predictions.

All implementations are available at https://github.com/Liangyushi/MiRGraph/tree/main.

## IV. Results

### A. Benchmark of miRTarBase MTI predictions

We compared RCGN [23], MRMTI [18], HGT [10], HGT BiLSTM, TransCNN and MiRGraph (i.e., HGT+TranCNN) in terms of the miRTarBase MTI predictions using standard evaluation metrics. In scenario 1, MiRGraph conferred the best prediction performance with the highest AUROC and AUPR (Table I). MiRGraph not only conferred AUROC and AUPR that are 1.44 and 2.14 percent higher than those from the runner-up method but also achieved the best performance on all other metrics (Table I). We also observed a large overlap among the three methods of HGT, TranCNN and MiRGraph in terms of the predicted MTIs, especially in true positive MTIs and true negative MTIs (Fig. 4). Nonetheless, we observed that the number of false positive MTIs predicted by MiRGraph was much smaller than that of HGT and TranCNN (Fig. 4(d)). The improved specificity is likely attributed to the fusion of heterogeneous network and sequence features. Scenario 2 is more challenging as it involves predicting MTIs on heldout miRNAs, whose groundtruth MTIs were not used as labels in during training (although all of the miRNAs are present in the HIN based TargetScan network). Nonetheless, MiRGraph also conferred the best prediction performance on most metrics (Table I).

**TABLE I.** Comparison of MTIs prediction performance on the miRTarBase MTIs in two training strategies scenarios. NP is the abbreviation of ‘Negative Precision’.

| Methods | AUPR | AUROC | Precision | Recall | F1 | Accuracy | FPR | Specificity | NP |
| --- | --- | --- | --- | --- | --- | --- | --- | --- | --- |
| Scenario 1 |  |  |  |  |  |  |  |  |  |
| RGCN | 0.7379 | 0.8369 | 0.6435 | 0.7073 | 0.6739 | 0.7764 | 0.1900 | 0.8100 | 0.8509 |
| HGT_BiLSTM | 0.7865 | 0.8612 | 0.7234 | 0.7049 | 0.7141 | 0.8156 | 0.1307 | 0.8693 | 0.8587 |
| HGT | 0.8418 | 0.9003 | 0.7422 | 0.7749 | 0.7582 | 0.8386 | 0.1305 | 0.8695 | 0.8885 |
| MRMTI | 0.8727 | 0.9198 | 0.7935 | 0.8014 | 0.7974 | 0.8671 | 0.1011 | 0.8989 | 0.9033 |
| MiRGraph(end-to-end) | 0.8955 | 0.9341 | 0.7893 | 0.8205 | 0.8046 | 0.8699 | 0.1062 | 0.8938 | 0.9113 |
| TransCNN | 0.9106 | 0.9418 | 0.8142 | 0.8398 | 0.8268 | 0.8851 | 0.0929 | 0.9071 | 0.9211 |
| MiRGraph(step-by-step) | **0.9320** | **0.9562** | **0.8569** | **0.8581** | **0.8575** | **0.9069** | **0.0695** | **0.9305** | **0.9311** |
| Scenario 2 |  |  |  |  |  |  |  |  |  |
| RGCN | 0.5249 | 0.6782 | 0.5067 | 0.4973 | 0.5020 | 0.7097 | 0.2018 | 0.7982 | 0.7921 |
| MRMTI | 0.5972 | 0.7475 | 0.5209 | 0.6610 | 0.5827 | 0.7214 | 0.2533 | 0.7467 | 0.8409 |
| HGT_BiLSTM | 0.6356 | 0.7810 | 0.8124 | 0.2164 | 0.3417 | 0.7548 | **0.0208** | **0.9792** | 0.7499 |
| HGT | 0.6574 | 0.7953 | 0.4214 | 0.8772 | 0.5693 | 0.6095 | 0.5021 | 0.4979 | **0.9068** |
| MiRGraph(end-to-end) | 0.7868 | 0.8785 | 0.6331 | 0.7867 | 0.7016 | 0.8031 | 0.1900 | 0.8100 | 0.9011 |
| TransCNN | 0.8057 | **0.8886** | 0.6526 | **0.7993** | 0.7185 | 0.8158 | 0.1773 | 0.8227 | 0.9077 |
| MiRGraph(step-by-step) | **0.8072** | 0.8862 | **0.7423** | 0.7065 | **0.7240** | **0.8415** | 0.1022 | 0.8978 | 0.8801 |

**Fig. 4.**
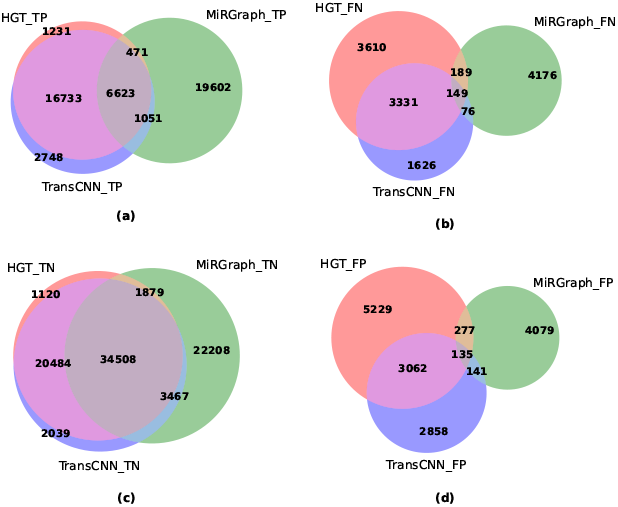
Venn diagram comparing HGT, TranCNN and MiRGraph predicted MTI. TP, TN, FP and FN stand for True positive, True Negative, False Positive, and False Negative, respectively.

In addition, the performance of RGCN is significantly worse than HGT, which demonstrates the advantage of the transformer layer in learning the attention weights between different relationships to better capture the heterogeneous information in the graph network. Interestingly, comparing HGT, RGCN, HGT BiLSTM and MRMTI, we found that the prediction performance of MRMTI (fusing graph and sequence information with RGCN and BiLSTM) is better than RGCN. On the other hand, HGT BiLSTM (i.e., using HGT and BiLSTM to learn graph and sequence information, respectively) is worse than HGT. This implies the limitation of the BiLSTM module compared to our TransCNN module given the same capability of graph feature representation. The ablation study also shows that the sequence features learned by the *Trans CNN* (AUROC 0.9418 and AUPR 0.9106) is more informative than the graph features learned by the *HGT* (AUROC 0.9003, AUPR 0.8418). Nonethless, combining the features learned by the two modules led to the highest performance, implying MiRGraph’s ability to learn complementary information from both sequences and HIN.

### B. Predicting log-fold change from miRNA transfections

We compared the ability of TargetNet [20] and miRGraph(step-by-step) in terms of the correlation between the level of expression down-regulation measured by *in-vitro* miRNA transfection experiments and the predicted LFCs. We ranked each miRNA-mRNA pair in the LFC dataset based on the predicted LFC scores, and examined the average of the LFC values of their expression among the top ranked predicted MTIs (Fig. 5 left panel). The top ranked targets by MiRGraph exhibit more negative LFC than the top ranked targets from TargetNet. The predicted LFC by MiRGraph also correlate better with the groundtruth LFC compared to TargetNet (Fig. 5 right panel).

**Fig. 5.**
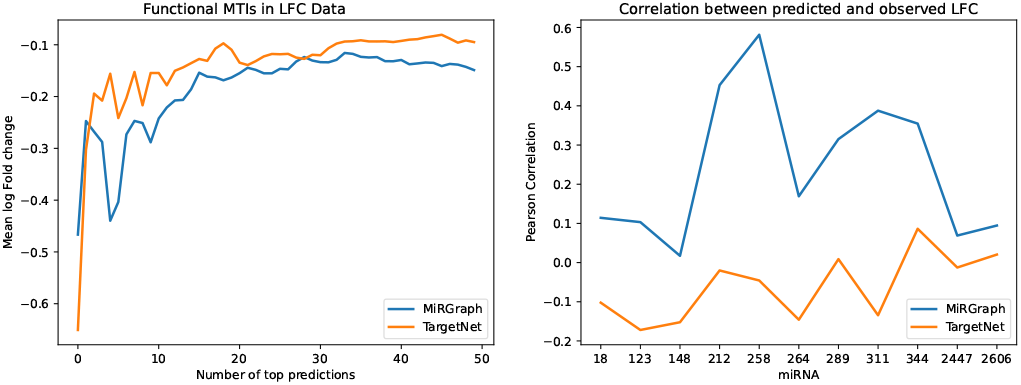
Predicting log-fold change (LFC) of gene expression from miRNA transfection experiments. (Left) Average LFC value of the top 50 targets from each of the 11 miRNAs. (Right) Pearson correlation coefficient between the predicted LFC and the groundtruth LFC values of the 11 miRNAs. The miRNAs indices [18, 123, 148, 212, 258, 264, 289, 311, 344, 2447, 2606] correspond to [‘hsa-miR-1-3p’, ‘hsa-miR-122-5p’, ‘hsa-miR-124-3p’,’hsa-miR-128-3p’, ‘hsa-miR-132-3p’, ‘hsa-miR-133a-3p’, ‘hsa-miR-142-3p’, ‘hsa-miR-148b-3p’, ‘hsa-miR-181a-5p’, ‘hsa-miR-7-5p’, ‘hsa-miR-9-5p’], respectively.

### C.Case study in breast cancer

We used MiRGraph trained on the miRTarBase dataset to investigate the target genes of an oncomir and miRNA regulators of an oncogene in cancer. In particular, we chose to investigate *hsa-miR-122-5p*, which is associated with cancer development and metastasis by targeting specific oncogenes [4]. Table II lists the top five MiRGraph-predicted target genes. Among them, *ABDH2* and *ACOT2* were supported by Tarbase v9.0 [24]. Then, we investigated the predicted miRNAs that bind to the oncogene *BRCA1*, which is a well-known hereditary gene associated with the risk of breast and ovarian cancers. Table II shows the top five predicted miRNAs and their supports. There are three out of five validated miRNAs i.e., *hsa-miR-124-3p, hsa-miR-16-5p, hsa-miR-21-5p*, which are implicated in tumorigenesis.

**TABLE II.**
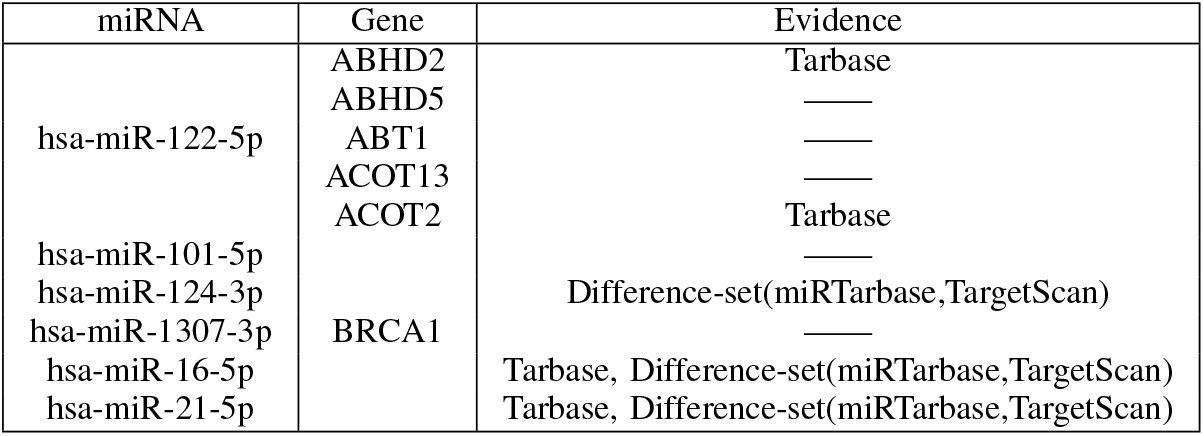
Top five candidate genes/miRNAs predicted by MiRGraph for hsa-miR-122-5p and BRCA1. Difference-set(miRTarbase,TargetScan) represents MTIs that exist only in miRTarbase and not in TargetScan, in other words, these data are not involoved in training and testing step.

| miRNA | Gene | Evidence |
| --- | --- | --- |
| hsa-miR-122-5p | ABHD2 | Tarbase |
|  | ABHD5 | — |
|  | ABT1 | — |
|  | ACOT13 | — |
|  | ACOT2 | Tarbase |
| hsa-miR-101-5p | BRCA1 | — |
| hsa-miR-124-3p |  | Difference-set(miRTarbase,TargetScan) |
| hsa-miR-1307-3p |  | — |
| hsa-miR-16-5p |  | Tarbase, Difference-set(miRTarbase,TargetScan) |
| hsa-miR-21-5p |  | Tarbase, Difference-set(miRTarbase,TargetScan) |

## V. Discussion

The investigation on the miRNA-target interactions can help reveal the complex gene regulation and disease treatment. Currently, few method can integrate multi-relationship network information and sequence information to identify functional MTIs. In this study, we develop MiRGraph, a multi-view deep learning framework to improve MTI prediction. Our experimental results suggest that MiRGraph conferred significant performance improvements in MTIs predictions over the state-of-the-art methods. Our case studies highlight putative targets of the oncomir *hsa-miR-122-5p* in cancer development and the putative miRNA regulators of *BRCA1*.

We consider several future extensions. MiRGraph is trained to distinguish whether the TargetScan-predicted MTIs are experimentally validated in miRTarBase. One caveat is that some TargetScan-predicted edges may be true positives but do not yet have experimental supports recorded in miRTarbase. We can use HITS-CLIP experimental data to identify negative edges. Furthermore, MiRGraph can be extended to utilize gene expression data to predict dynamic MTIs.

## Acknowledgments

This work is supported by the National Natural Science Foundation of China (#62032007, #62372165 and #62202152) and the Natural Sciences and Engineering Research Council (NSERC) Discovery Grant (RGPIN-2019-0621).

